# The microbiota of wooden cheese ripening boards is a rich source of antimicrobial-producing bacteria against *Listeria monocytogenes*

**DOI:** 10.1101/2024.11.14.623661

**Authors:** Yuxing Chen, Ibrahim Zuniga Chaves, Garret Suen, TuAnh N. Huynh

**Affiliations:** Food Science Department, University of Wisconsin – Madison; Department of Bacteriology, University of Wisconsin - Madison

## Abstract

Wooden boards are essential tools in cheese ripening and there are accumulating observations suggesting their antimicrobial effect against foodborne bacterial pathogens, such as *Listeria monocytogenes*. However, poor bacterial recovery of bacteria from wood can confound quantification of pathogen burdens. To assess *L. monocytogenes* survival on wooden cheese boards, we applied a disruptive grinding method and tracked native board-associated bacterial counts as controls. Our data revealed that *L. monocytogenes* declines on clean zones of wooden boards, but can replicate on areas where there is suitable cheese. Our microbiota analysis revealed diverse bacterial communities on wooden board surfaces, with a prominent presence of *Brevibacterium*, *Brachybacterium*, and *Staphylococcus* genera. We further identified seven bacterial species that inhibit *L. monocytogenes*, belonging to *Bacillus*, *Staphylococcus*, and *Serratia* phyla, as well as *Lactococcus lactis*. We focused on a *Bacillus safensis* isolate as a novel biocontrol agent candidate, and found it to potently inhibit *L. monocytogenes* via secreted antimicrobial factors. Our genomic, bioinformatic, and biochemical analyses indicate that those factors are likely antimicrobial peptides encoded by multiple biosynthetic gene clusters, several of which are unique to *B. safensis* and have not been characterized. A sub-inhibitory concentration of *B. safensis* supernatant induces a significant down-regulation of prophage elements and up-regulation antimicrobial stress response in *L. monocytogenes*. Taken together, our findings indicate that the wooden board microbiota is a rich source of antimicrobial-producing bacteria with potential applications in foodborne pathogen control strategies.

**IMPORTANCE:** Despite stringent food safety measures, *L. monocytogenes* foodborne outbreaks remain frequent with high hospitalization and mortality rates. Removal of *L. monocytogenes* from food processing environments is extremely challenging, because this pathogen is ubiquitous and encodes a wide array of stress response mechanisms that enable it to thrive under harsh conditions. Our study found that clean wooden boards used in cheese ripening inhibit *L. monocytogenes*, causing a noticeable decline in pathogen population following surface inoculation. Bacterial communities on wooden cheese boards are rich and diverse, and harbor many species that produce antimicrobial compounds against *L. monocytogenes*, with the example of a new *B. safensis* isolate. Therefore, wooden board microbiota is a promising source for future antimicrobial discovery efforts.

## INTRODUCTION

Wooden boards have long been used for cheese ripening and remain popular among artisanal cheese makers, accounting for the ripening of ∼500,000 tons of cheese in Europe every year (1). The porous and hygroscopic properties of wood are considered essential in the development of cheese sensory quality during ripening. Indeed, cheese made with wooden tools has been reported to exhibit a richer volatile compound profile than cheese made with other tools (2). Wood used in cheese ripening is typically dried to 15-18% humidity and without chemical treatments. Wood with high humidity might support mold or *Pseudomonas fluorescens*. By contrast, dry wood might promote the development of *Serratia* species, which cause thick, strong rinds, and red defects (1). Therefore, cheese makers often intentionally select woods with suitable hydrometry to achieve desirable levels of cheese humidity and ripening kinetics.

Wooden boards harbor a surface microbiota that is rich in composition and diverse among boards (1, 3–6). For instance, a comprehensive survey of wooden boards in 18 dairy facilities in Sicily, Italy revealed at least 43 bacterial genera on each board (5). Although those boards exhibited distinct microbiota compositions, some bacterial groups are commonly present in significant abundances, including *Staphylococcus*, *Brevibacterium*, and *Corynebacterium*. Interestingly, these three genera were also found to be dominant in a survey of wooden cheese boards in Wisconsin, United States (4), suggesting that these bacteria are among core members of cheese board microbial communities.

Bacteria on wooden board surfaces are considered a major source of cheese rind microbiota whose metabolic activities drive cheese flavor development (1, 7). The biofilm formed by these bacteria is stable and can withstand some sanitation treatments. This persistence has raised food safety concerns, highlighted by a brief FDA ban of wooden boards in cheese ripening. Although the ban was quickly reversed, and wooden boards have a long history of safe use, dairy processing facilities are frequently contaminated with foodborne pathogens, thorough assessments of pathogen behavior on cheese boards are necessary to guide ripening process design.

*Listeria monocytogenes* is an invasive foodborne pathogen that is frequently associated with dairy product outbreaks. *L. monocytogenes* isolates from dairy products are particularly well-adapted to host conditions and gut colonization. A majority of those isolates are of clonal complex 1, which is hyper-virulent and strongly associated with human clinical samples (8). *L. monocytogenes* is ubiquitously present in dairy farms and prevalently shed by cattle, and therefore easily contaminates raw milk and dairy processing plants (9–12). Although pasteurization is effective at killing *L. monocytogenes*, this pathogen can contaminate dairy products at any processing steps following pasteurization (13). The persistence of *L. monocytogenes* in dairy processing plants is well documented, and this pathogen can be detected at many locations within a plant, especially in difficult-to-reach areas such as drains, cracked surfaces, joints of equipment and conveyor belts (13). From a physiological standpoint, *L. monocytogenes* has an array of stress response mechanisms that enable its persistence under dairy processing conditions and replication in dairy products. For instance, the general stress response regulator, σ^B^, is activated by many stress conditions and upregulates more than 200 genes to adapt to low pH, high osmolarity, and refrigeration, all of which are relevant in dairy production (14). Soft cheeses are particularly permissive to *L. monocytogenes* replication due to comparatively high humidity and neutral pH. Furthermore, *L. monocytogenes* is notorious for robust biofilm formation on different materials, and these biofilms provide additional protection against antimicrobial treatments (15, 16).

Despite the widespread presence of *L. monocytogenes* in dairy processing plants, wooden boards have not been identified as the source of *L. monocytogenes* contamination in cheese. On the contrary, there is evidence for the antimicrobial effect of wooden boards. For instance, wood transfers significantly less *L. monocytogenes* to cheese than plastic and glass (17). Furthermore, *L. monocytogenes* burdens have been shown to significantly reduce following inoculation on native wooden boards (18). Interestingly, heat-inactivated wooden boards allows *L. monocytogenes* replication, suggesting that the board surface microbiota confers inhibition (18). However, since *L. monocytogenes* recovery rate from wood is poor, at no more than 30% even with destructive methods (19), alternative experimental designs are necessary to accurately quantify *L. monocytogenes* survival rates on wooden boards. Nevertheless, a recent study found that culturable bacteria from wooden boards inhibit *L. monocytogenes* in laboratory culture, although antagonistic bacteria have not been systematically identified (20).

In this study, we aimed to assess *L. monocytogenes* survival on wooden cheese board surface, and to systematically identify board-associated bacteria that inhibit *L. monocytogenes*. To address the technical challenge of poor bacterial recovery from wood, we concurrently tracked both *L. monocytogenes* and wooden board-associated bacteria as controls. We found *L. monocytogenes* burdens to decline in most instances on washed and native boards, but at different rates and extents among boards. In one exception, we observed *L. monocytogenes* replication relative to board-associated bacteria, although cheese residues likely contributed to this replication. Our survey of a small set of wooden boards from three cheese makers revealed rich microbiota compositions with the prominence of *Staphylococcus*, *Brevibacterium*, and *Brachybacterium* phyla, consistent with two previous surveys of wooden boards elsewhere (4, 5). Exploiting this microbial diversity, we identified six bacterial species of the *Bacillus*, *Staphylococcus*, *Lactococcus*, and *Serratia* genera that inhibit *L. monocytogenes*. We focused on a *Bacillus safensis* isolate as a novel candidate for biocontrol agents, and found it to potently inhibit *L. monocytogenes*, likely through secreted antimicrobial peptides. In response, *L. monocytogenes* significantly down-regulated the prophage and monocin elements. Together, our findings indicate that wooden board microbiota is a rich source of natural antimicrobials against *L. monocytogenes*.

## RESULTS

### Wooden boards harbor a diverse microbiota

Despite long research interests in the microbial ecology of wooden cheese boards, only two studies have systematically determined the wooden board microbiota by next-generation sequencing (4, 5). To expand our knowledge base of microbial diversity on wooden cheese boards, we obtained seven wooden boards from three different cheese makers in Wisconsin, United States, including native and washed boards from each cheese maker. These cheese makers are distinct from those surveyed in a previous study (4). We noticed that each board had a circular cheese mark where a cheese block had been placed during ripening, as mapped out in **Fig. 1**. We therefore analyzed the bacterial compositions of “clean” zones separately from “cheese” zones on each board to avoid the confounding presence of cheese microbiota in our analyses.

**Figure 1:**
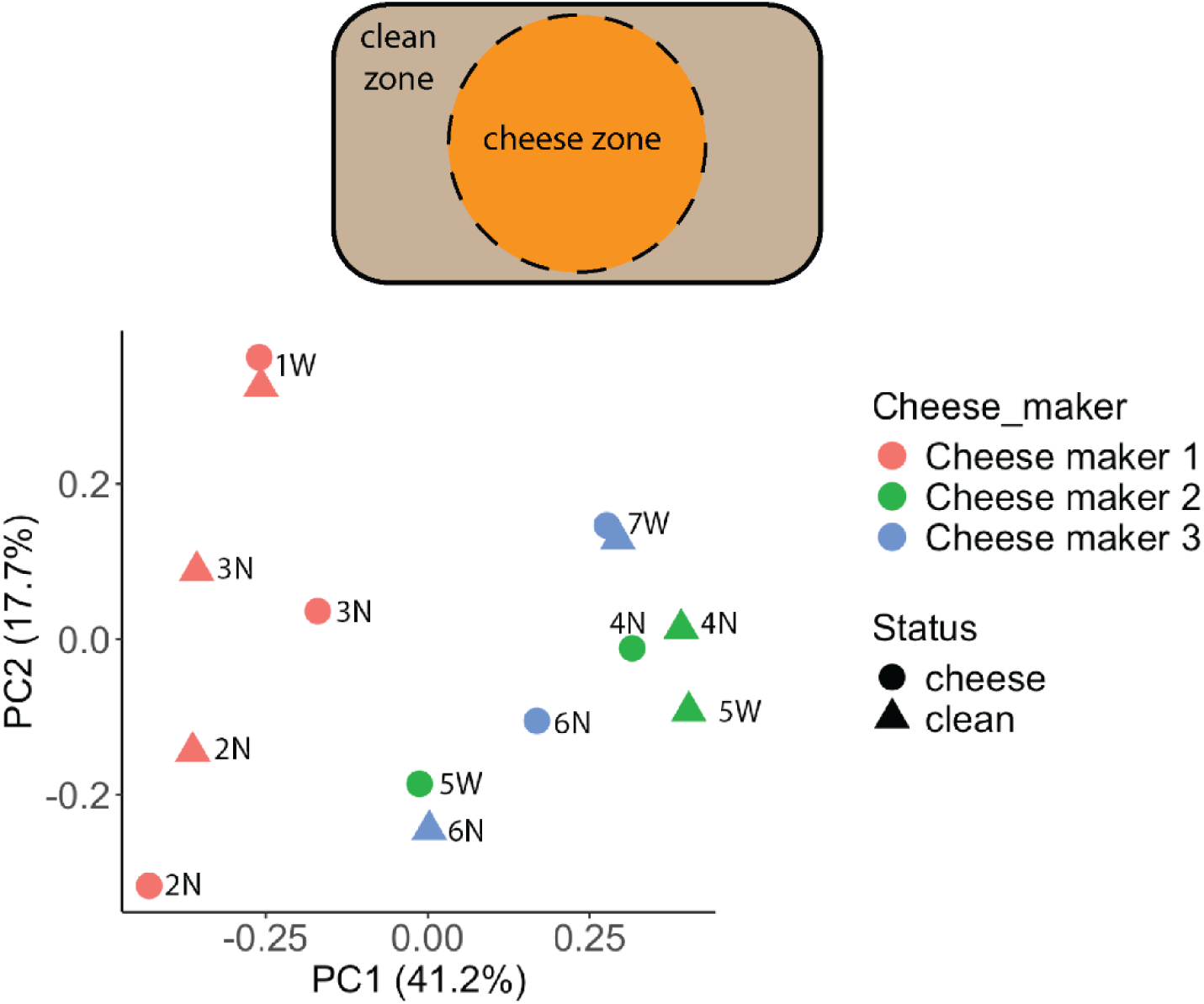
Wooden boards harbor distinct microbiota compositions. Top panel: clean zone and cheese zone on each wooden board. Bottom panel: Principle component analysis of bacterial communities on wooden boards from three cheese makers. Each data point represents the microbiota composition of a native or washed wooden board from a cheese maker. Each board is marked with board number, N indicates native boards, W indicates washed boards.

Total DNA was extracted from the top 2 mm of each board, obtained by grinding. After denoising and decontamination, we obtained a total of 183918 high-quality reads and grouped them into 62 operational taxonomy units (OTUs). These OTUs belong to 12 phyla, 16 classes, 44 orders, 72 families, and 93 genera. Due to the limited availability of boards, we did not obtain sufficient biological replicates of each board type to reliably compute α-diversity indices. However, we noticed rich and diverse microbiota, with each board harboring between 8-58 genera (median of 22).

As expected, bacterial compositions were distinguishable for wooden boards from different cheese makers, although those from cheese maker 1 was the most dissimilar (**Fig. 1**). From each cheese maker, microbiota compositions were also distinguishable between native and washed boards, with most notable changes in the abundances of *Staphylococcus* and *Virgibacillus*. Overall, the most three abundant bacteria across all boards belonged to three genera: *Staphylococcus* (from 0.06% to 52%, median 17%), *Brevibacterium* (from 4% to 69%, median 30%), and *Brachybacterium* (from 0.4% to 32%, median 3.5%), consistent with previous surveys of cheese board and cheese rind microbiota compositions (4, 5, 21) (**Fig. 2**).

**Figure 2:**
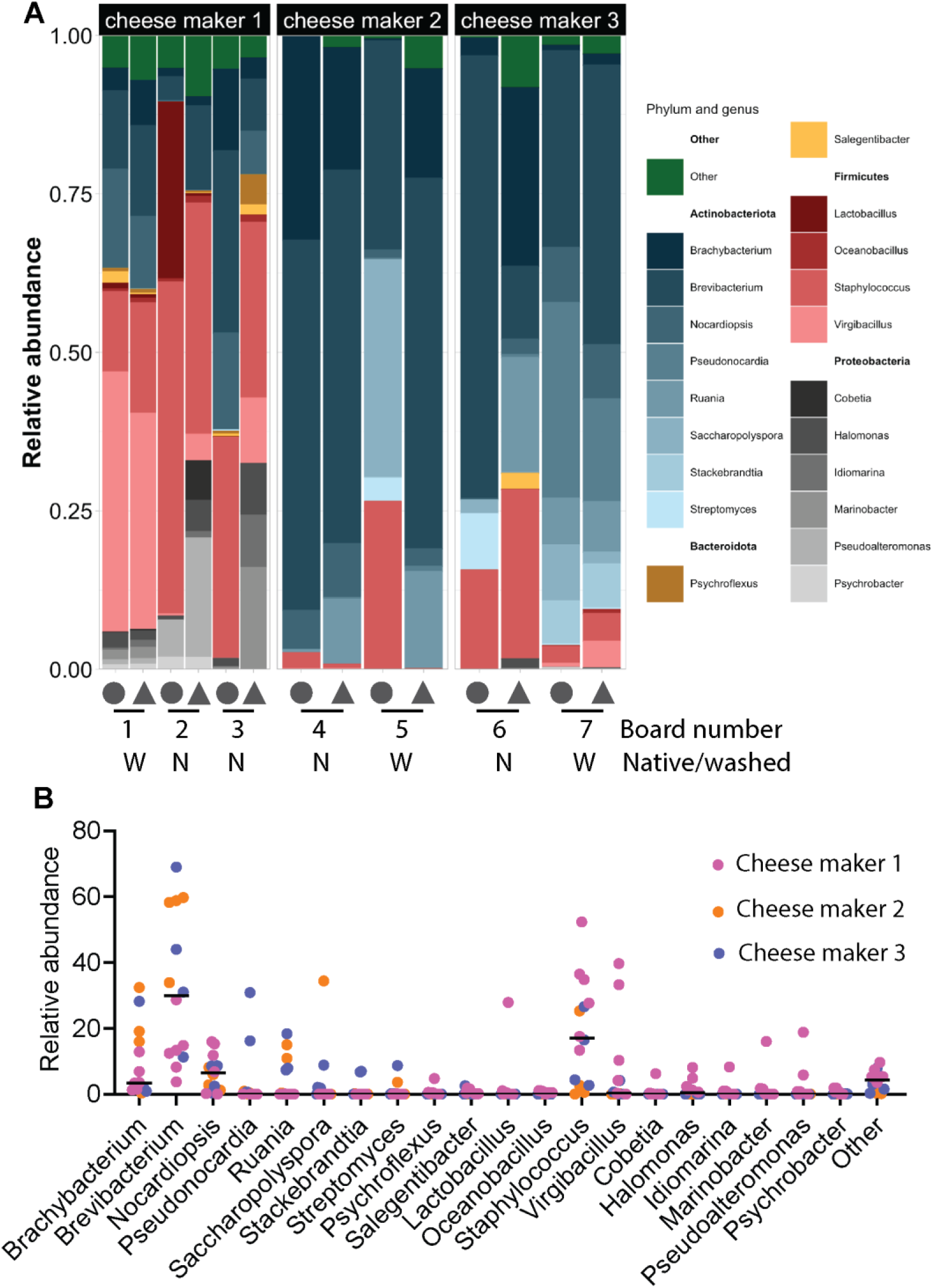
Bacterial communities on wooden boards are diverse. Relative abundances of the twenty most abundant phyla on seven wooden boards are organized for each board (**A**) or each phylum (**B**). In panel A, circle symbol indicates cheese zone, and triangle indicates clean zone on each board. Bars in panel B are median abundances.

While sharing some common bacteria, each board in our microbiota survey exhibited a unique composition (**Fig. 2**). Two native boards from cheese maker 1 were dominated by *Staphylococcus* (13% and 52%), *Brachybacterium* (1% and 13%), and *Brevibacterium* (4% and 29%). We also noticed that cheese blocks from native board 2 had a high abundance of *Lactobacillus* (∼28%), likely transferred from cheese. Somewhat distinct from others, board 3 was also abundant in *Nocardiopsis* (7-10%). By comparison, the washed board from this cheese maker (board 1) was significantly reduced in *Staphylococcus* and substantially increased in *Nocardiopsis*.

Boards from cheese maker 2 (boards 4 and 5) were dominated by *Brevibacterium* (34-60%) and *Brachybacterium* (16-32%), with an additional abundance of *Nocardiopsis* (1.2-8%). Additionally, *Ruania* was abundant on clean blocks from this cheese maker (11-15%), but not on cheese blocks, suggesting that these bacteria are of wood or environmental origins.

Wooden boards from cheese maker 3 (boards 6 and 7) were overall abundant with *Brachybacterium* (1-28%), *Brevibacterium* (11-69%), and *Staphylococcus* (3-27%). As observed in cheese maker 1, *Staphylococcus* was much reduced in abundance on a washed board than on a native board (∼3% compared to ∼20%, respectively). By contrast, *Nocardia* and *Pseudonocardia* were much more abundant on a washed board from this cheese maker (∼8% and ∼23%, respectively).

### *L. monocytogenes* declines on the clean zones of most wooden cheese boards

A previous study showed *L. monocytogenes* to be killed on the surface of native wooden boards (18). However, recovery of *L. monocytogenes* from wood is notoriously poor, and could confound quantification of *L. monocytogenes* burdens on wood under prolonged incubation. We therefore examined both *L. monocytogenes* burdens and native cheese board bacteria as controls for our recovery method and incubation condition.

We obtained four boards from cheese maker 1, including two native boards similar to boards 2 and 3, and two washed boards similar to board 1 in our microbiota analyses. We again separated clean blocks from those inside the cheese zone, because cheese residues can impact *L. monocytogenes* growth and survival (**Fig. 3A**). Each board was cut into square blocks, surface inoculated with *L. monocytogenes*, and incubated at 12°C, 85% relative humidity for 20 days. To recover bacteria from those blocks, we applied a destructive method, shown to have the highest recovery rate, which involved shaving, grinding, and rigorous suspension in phosphate-buffered saline.

**Figure 3:**
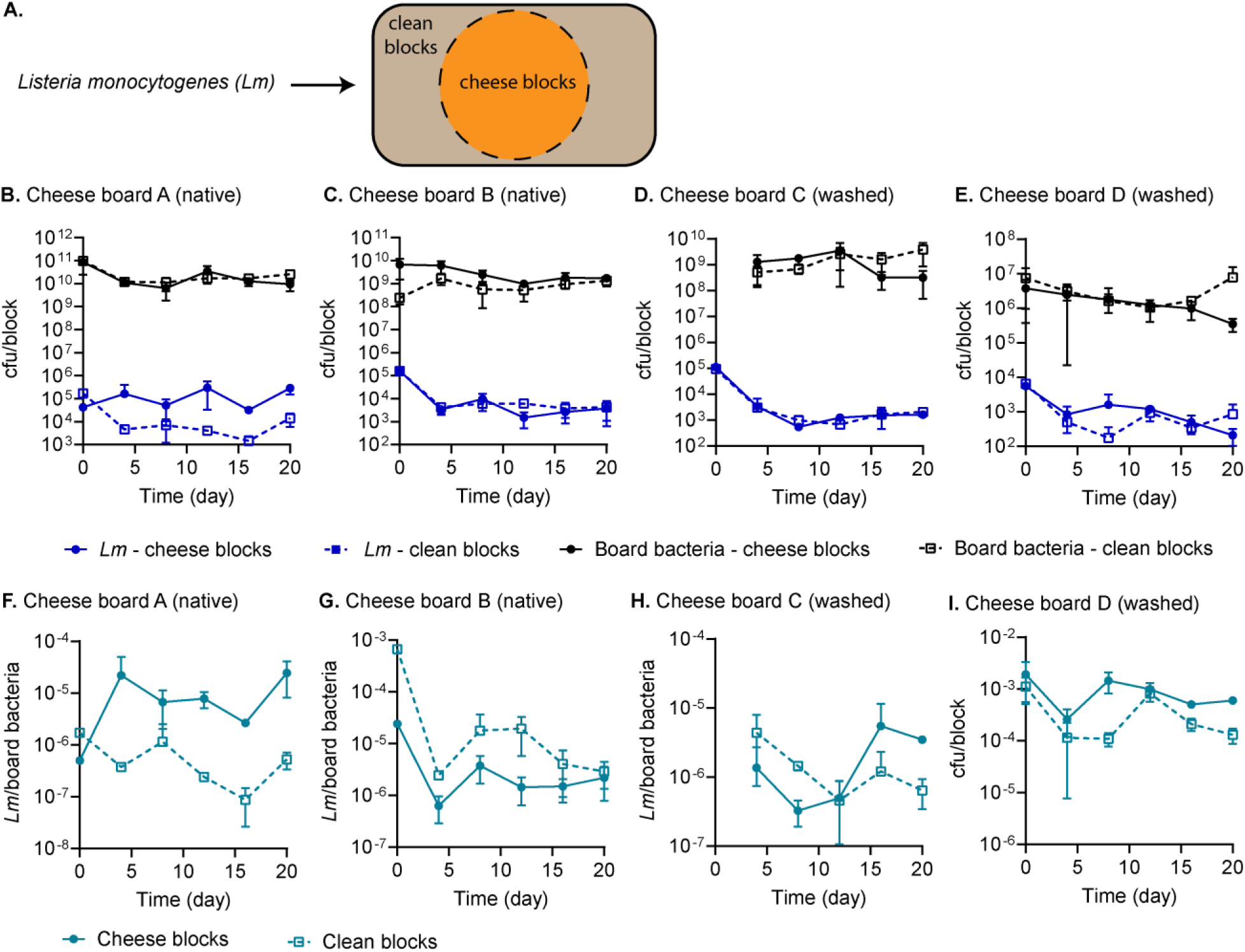
*L. monocytogenes* generally declines on wooden cheese boards following surface inoculation. **A.** Schematic diagram of a wooden board with clean and cheese blocks, which were separated prior to surface inoculation with *L. monocytogenes*. **B-E.** Burdens of *L. monocytogenes* and board-associated bacteria on each block. **F-I.** Relative burdens of *L. monocytogenes*, normalized to those of board-associated bacteria from each block. Native board-associated bacteria were not collected from board C at time point 0.

Over 20 days, board-associated bacterial counts remained steady (**Fig. 3B-E**). The native boards harbored ∼ 10^9^-10^10^ cfu/block (∼ 1.7×10^7^-1.7×10^8^cfu/cm^2^), whereas the washed boards harbored ∼ 2×10^6^ – 10^9^ cfu/block (∼ 1.7×10^4^ – 1.7×10^7^ cfu/cm^2^). Examining raw counts of *L. monocytogenes* on wooden blocks, we found them to generally decline over the incubation period. The reduction in *L. monocytogenes* was most evident on clean blocks, with a ∼ 1.5-log cfu decrease over the first four days, and a similar magnitude of reduction was also apparent on cheese blocks of boards B, C, D. The only exception to this trend was on cheese blocks of board A, where *L. monocytogenes* burdens exhibited a modest increase on day 20 compared to day 0 (**Fig. 3B-E**).

Upon normalizing *L. monocytogenes* counts to those of board-associated bacteria on each block, we observed an apparent replication of *L. monocytogenes* on cheese blocks of board A, with a nearly 2-log increase in cfu counts between days 0 and 20 (**Fig. 3F-I**). However, *L. monocytogenes* burdens otherwise declined on cheese blocks of other boards, and on clean blocks of all boards. Clean blocks of board B were the most inhibitory, causing a ∼2-log cfu reduction in *L. monocytogenes* over 20 days. The magnitude of *L. monocytogenes* decline on other blocks was ∼ 0.5-1-log cfu.

### Wooden board microbiota harbor bacteria that inhibit *L. monocytogenes*

A previous study identified *Leuconostoc mesenteroides* and *Staphyloccus equorum* from wooden boards to inhibit *L. monocytogenes* (20). The microbial diversity of wooden boards in our study suggests that there might be more bacterial species from these boards that antagonize *L. monocytogenes*. Therefore, we systematically isolated cheese board-associated bacteria by recovering wood chip suspensions in different growth media, including Tryptic Soy agar, Brain Heart Infusion agar, MRS, and PCAMS (22). This effort yielded approximately 500 bacterial isolates that were screened for antimicrobial activity by spotting on a *L. monocytogenes* lawn on suitable agar. Around 220 isolates produced a zone of clearance, indicating antimicrobial activity, and these were further purified through multiple rounds.

Through the above culture-based method, we obtained 54 bacterial isolates representing four major types of colony morphology. Identification by a MALDI-TOF Biotyper revealed that those isolates belong to *Bacillus*, *Staphylococcus*, *Lactococcus*, and *Serratia* genera, although the species could not be unambiguously identified. Therefore, we selected 11 isolates for 16S rRNA sequencing, and identified them as listed in **Table 1**. These bacteria include two *Bacillus* species, two *Staphylococcus* species, *Lactococcus lactis*, and *Serratia marcescens*. Three of these species have been characterized for *L. monocytogenes* inhibition: *L. lactis* produces bactericidal bacteriocins (23); *Staphylococcus equorum* secretes micrococcin P1 that is bacteriostatic (20, 24); and *S. marcescens* produces a serrawettin lipopeptide that is bactericidal (25).

**Table 1:**
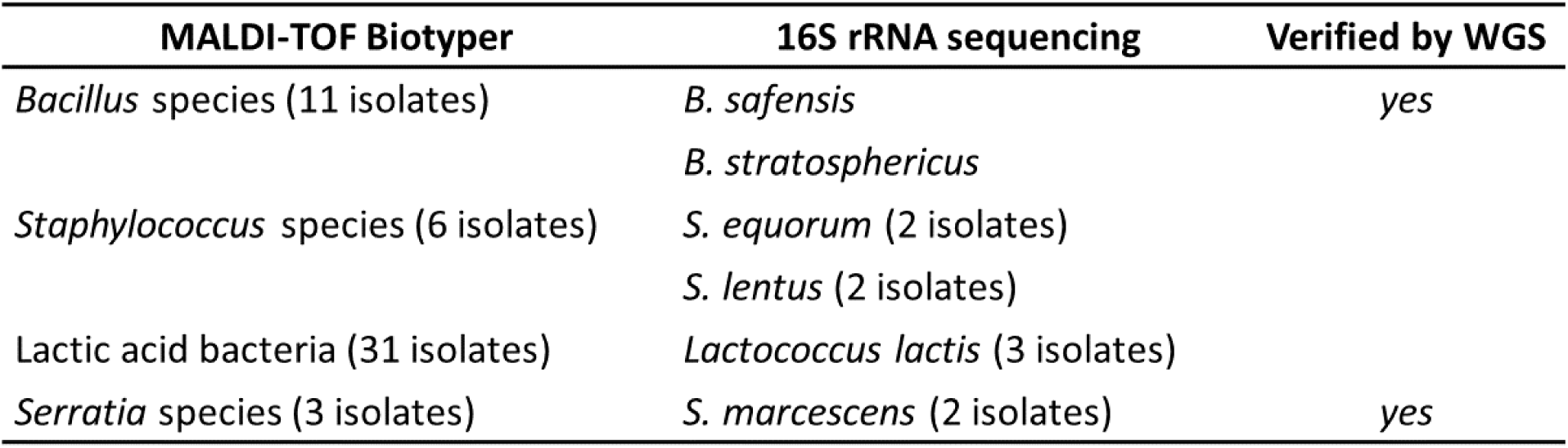
Identification of antimicrobial-producing bacteria from wooden cheese boards.

### Bacillus safensis secretes antimicrobials against L. monocytogenes

*B. safensis* and *B. stratosphericus* had not been characterized for antimicrobial activities against *L. monocytogenes*, and both were potent and consistent inhibitors of *L. monocytogenes* in our screen (**Fig. S1**). We chose to focus on *B. safensis*, an environmental bacterium of soil origins with antifungal activity (26).

The zone of *L. monocytogenes* clearance surrounding *B. safensis* growth suggests that *B. safensis* might produce and secrete antimicrobials against *L. monocytogenes*. To further test this possibility, we plated serial dilutions of *L. monocytogenes* away from *B. safensis* on an agar medium. *L. monocytogenes* was indeed inhibited by *B. safensis* despite spatial separation, confirming that *B. safensis* secretes anti-*Listeria* factors (**Fig. 4A**). In this experimental set up, antimicrobial synthesis and secretion were somewhat delayed, since *L. monocytogenes* was only inhibited if *B. safensis* was pre-grown for at least one day prior to *L. monocytogenes* plating. The inhibitory effect was diminished by 4-day old *B. safensis*, suggesting that secreted antimicrobials were unstable by this time point. Together, these data also reveal that *B. safensis*-derived antimicrobials accumulate as the culture grows and their synthesis is not triggered by *L. monocytogenes*.

**Figure 4:**
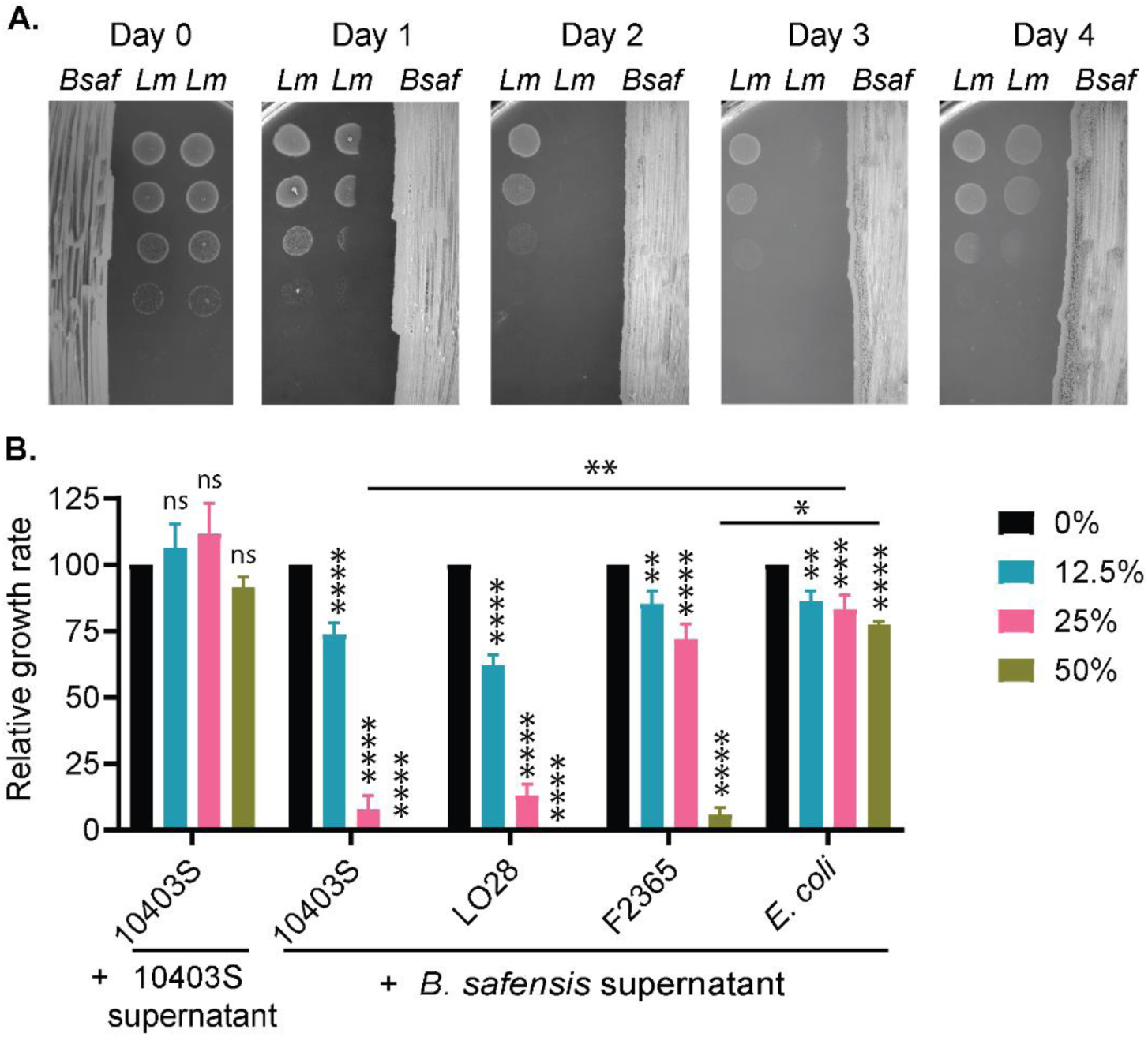
*B. safensis* inhibits *L. monocytogenes* growth via secreted antimicrobial factors. **A.** Inhibition of *L. monocytogenes* growth on BHI agar. *B. safensis* was streaked on one half of the agar and pre-grown for 0 – 3 days at 30°C. Ten-fold dilutions of *L. monocytogenes* cultures were spotted on the other half of the agar, on indicated number of days after *B. safensis* was streaked. **B.** Inhibition of *L. moncytogenes* growth in BHI broth at varying concentrations of *B. safensis* supernatant. Relative growth rate was calculated as growth rate in the presence of added supernatant, normalized to that in BHI broth only. Statistics: one-way ANOVA within each strain, and two-way ANOVA among strains. ns: non-significant; **, P < 0.01; ****, P < 0.0001.

To evaluate secreted antimicrobials in liquid cultures, we obtained a cell-free culture supernatant from *B. safensis* that had been grown to a high density (OD_600_ >10), and tested it against *L. monocytogenes* growth. As controls, we also obtained *L. monocytogenes* culture supernatants from the same culture condition. We found *L. monocytogenes* to be potently inhibited by supernatant from *B. safensis*, but not *L. monocytogenes* (**Fig. 4B**). Furthermore, *E. coli* MG1655 was not inhibited by *B. safensis* supernatant. This observation indicates that *L. monocytogenes* inhibition likely occurs through antimicrobial activity of *B. safensis* rather than nutrient depletion in spent media.

Our assessments of *L. monocytogenes* inhibition thus far had focused on strain 10403S (sequence type 85). Because *L. monocytogenes* is genetically diverse, we expanded our analysis to include other commonly studied strains: F2365 (sequence type 1), EGD (sequence type 12), EGD-e (sequence type 35), and LO28 (sequence type 210) (27, 28). Strain F2365 was the most resistant, but all strains were completely inhibited by 50% *B. safensis* supernatant (**Fig. 4B**).

### Cheese board-associated *B. safensis* encodes many biosynthetic gene clusters with predicted antimicrobial functions

As a first step towards developing *B. safensis* for antimicrobial production towards *L. monocytogenes*, we performed whole genome sequencing to gain genetic knowledge for the isolate obtained in this study. Our cheese board *B. safensis* isolate, hereafter called CB375, has a genome of approximately 3.8 Mbp with 41.64% GC content, harboring 3,763 genes, encoding 3,692 proteins, 64 tRNA genes and 6 rRNA genes (**Fig. 5A**). Of the 3,781 CDS predicted, 1,184 are proteins of unknown function, and 2,579 proteins have functional assignments, including 1,312 proteins with Enzyme Commission (EC) numbers, 1,799 with Gene Ontology (GO) assignments and 752 proteins that can be mapped to Kyoto Encyclopedia of Genes and Genomes (KEGG) pathways. By phylogenetic and average nucleotide identity (ANI) analyses, we confirmed that CB375 is *B. safensis*, and most closely related to strain FUA2118 (**Fig. S2** and **Table S1**). Antimicrobial resistance gene detection by the CARD database (29) indicated that *B. safensis* CB375 harbors a putative chloramphenicol acetyltransferase (CAT), that exhibits 42% identity and 66% similarity to CAT from commonly used bacterial expression plasmids (**Fig. S3**). In LB broth, *B. safensis* CB375 was noticeably more chloramphenicol resistant than *B. subtilis* type strain 168 that does not carry a *cat* gene, although their minimum inhibitory concentrations were similar: approximately 5 μg/mL for *B. safensis* and 4.5 μg/mL for *B. subtilis* (**Fig. 5B**).

**Figure 5:**
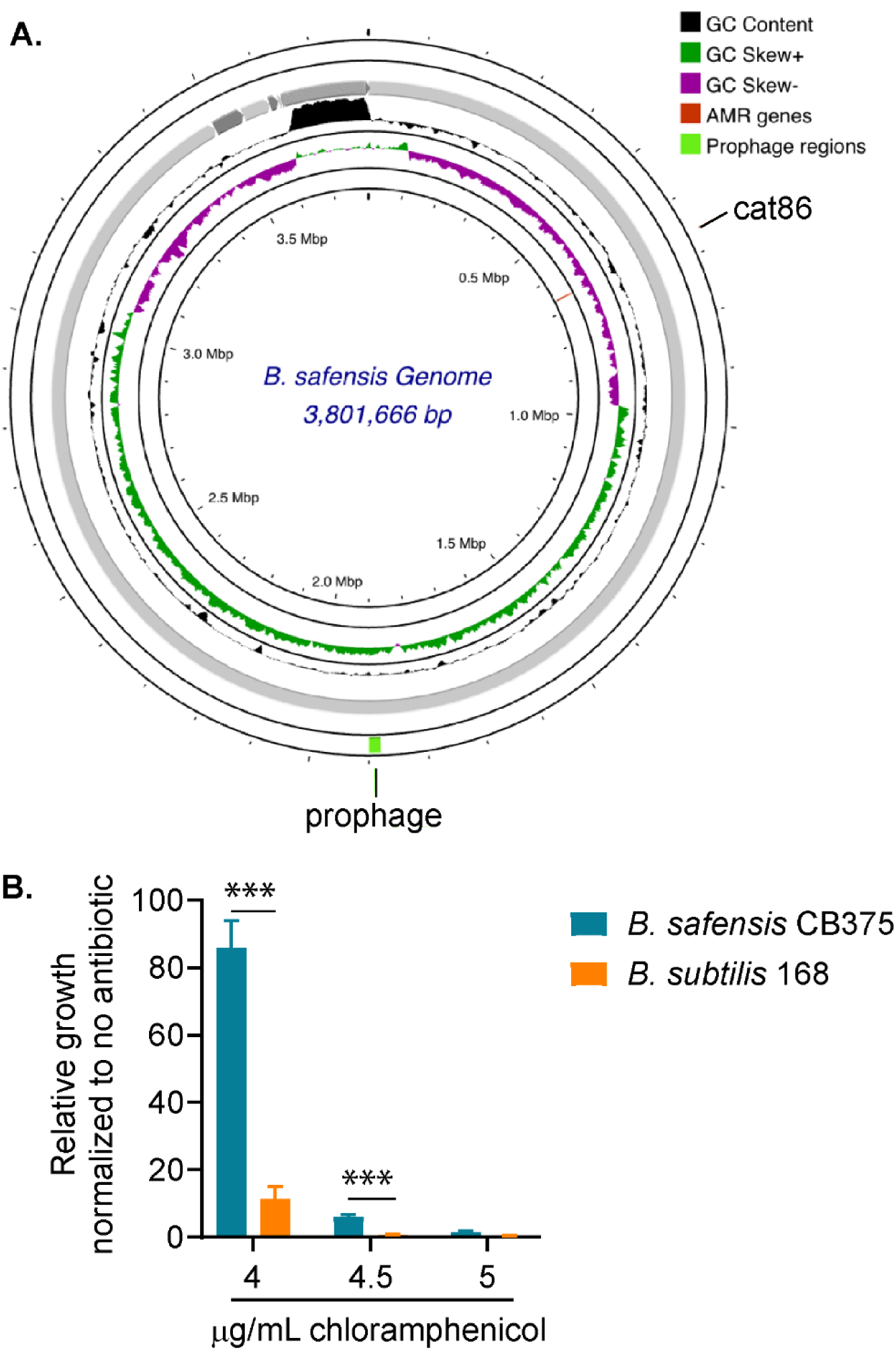
Genomic analysis of *B. safensis* strain CB375. **A.** The assembled *B. safensis* CB375 genome with a predicted chloramphenicol acetyltransferase gene (*cat86*). **B.** Chloramphenicol resistance of *B. safensis* CB375 and *B. subtilis* 168, grown in LB at 37°C. For each bacterium, growth at each chloramphenicol concentration was normalized to growth in LB only. Statistical analysis was performed by Student’s for the indicated pairs: ***, P < 0.001.

*Bacillus* species produce a wide range of antimicrobial peptides, encoded by biosynthetic gene clusters (BGCs). We therefore mined the *B. safensis* CB375 genome for BGCs using the antiSMASH database. This analysis revealed 14 BGCs, most of which are predicted to encode non-ribosomal peptide synthases (NPRS) (**Table 2**). Four BGCs within *B. safensis* CB375 are highly similar to those producing known compounds. BGC5 perfectly matches with the reference BGC for bacillibactin, a siderophore that has antibacterial and antifungal activities due to iron sequestration (30). BGC2 is 91% similar to plantazolicin, a peptide of the RiPP family (ribosomally synthesized and post-translationally modified peptides) with narrow spectrum activity towards *B. anthracis*, *B. cereus*, and *B. thurigiensis* (31). BGC6 and BGC12 exhibit 85% similarity to bacilysin and sporulation killing factor, respectively (32–35). Five BGCs have low similarity to fengycin (an antifungal peptide), schizokinen (a siderophore), lichenysin, and sporulation killing factor (36, 37). Of note, although BGC11 and BGC14 are both predicted to produce lichenysin, their sequences are distinct and most likely synthesize different compounds. The remaining five BGCs do not match with reference BGCs, and are unique to *B. safensis* and closely related species (**Table 2**).

**Table 2:** Predicted biosynthetic gene clusters within *B. safensis* CB735.

| BGC no. | Annotation | Similar known BGC | Reference organism |
| --- | --- | --- | --- |
| 1 | Betalactone | fengycin<br>53% similarity to BGC0001095 | *B. velezensis* |
| 2 | LAP;RRE-containing | plantazolicin<br>91% similarity to BGC0001173 | *B. pumilus* |
| 3 | NI-siderophore;terpene | schizokinen<br>62% similarity to BGC0002633 | *Leptolyngbya sp.* |
| 4 | RRE-containing | No matches found |  |
| 5 | NRP-metallophore;NRPS | bacillibactin<br>100% similarity to BGC0000309 | *B. subtilis* |
| 6 | other | bacilysin<br>85% similarity to BGC0001184 | *B. velezensis* |
| 7 | betalactone | No matches found |  |
| 8 | RiPP-like | No matches found |  |
| 9 | T3PKS | No matches found |  |
| 10 | Terpene | No matches found |  |
| 11 | NRPS | lichenysin<br>50% similarity to BGC0000381 | *B.licheniformis* |
| 12 | Sactipeptide | sporulation killing factor<br>85% similarity to BGC0000601 | *B. subtilis* |
| 13 | NRPS | surfactin<br>43% similarity to BGC0000433 | *B. velezensis* |
| 14 | NRPS | lichenysin<br>14% similarity to BGC0000381 | *B. licheniformis* |

### *B. safensis*-derived antimicrobials that inhibit *L. monocytogenes* are peptides or proteins

Given the presence of many antimicrobial peptide-encoding BGCs within the *B. safensis* genome, we sought to isolate secreted peptides from *B. safensis* by ammonium sulfate precipitation of its culture supernatant. These concentrated preparations of secreted proteins and peptides were bactericidal towards *L. monocytogenes* in a dose-dependent manner, with up to ∼6 log cfu of killing after 1 hour of exposure (**Fig. 6A**). Furthermore, proteinase K treatment abolished *L. monocytogenes* inhibition, further verifying that antimicrobial compounds are proteins or peptides in nature (**Fig. 6B**). Combined with the predicted presence of multiple BGCs, *B. safensis* CB375 most likely inhibits *L. monocytogenes* through secreted antimicrobial peptides.

**Figure 6:**
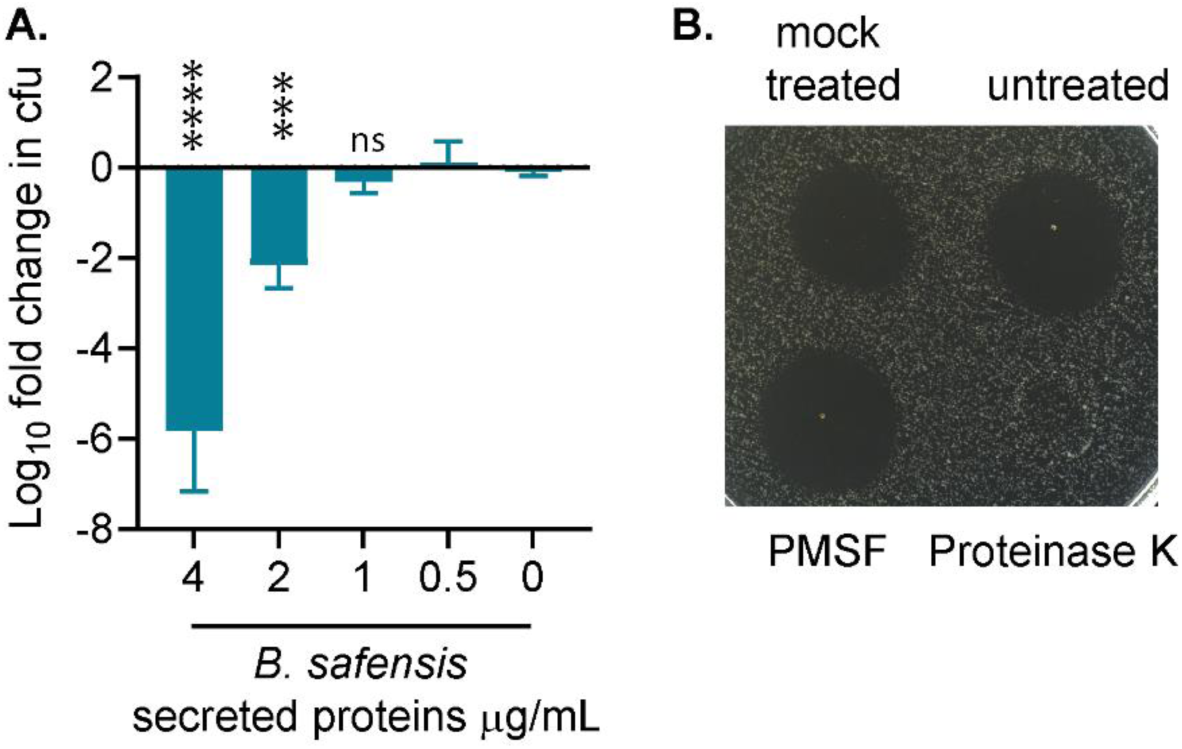
*B. safensis*-secreted antimicrobials against *L. monocytogenes* are proteins or peptides. **A.** Killing of *L. moncytogenes* by concentrated proteins and peptides from *B. safensis* culture supernatants. *B. safensis* supernatants were concentrated with ammonium sulfate and resuspended in PBS, then used to treat *L. monocytogenes* cultures, at approximately 10^9^ cfu/mL, for an hour. Statistical analysis was performed by one-way ANOVA, comparing each treatment with the control. ****, P < 0.0001; ***, P < 0.001, ns: non-significant. **B.** Concentrated *B. safensis* supernatants were treated with proteinase K for 2 hours at 37°C, 0.5mM PMSF (phenylmethylsulfonyl fluoride), or mock-treated, and spotted on a *L. monocytogenes* lawn to inspect for antimicrobial activities.

### *B. safensis* supernatant induces specific a stress response by *L. monocytogenes*

Although *L. monocytogenes* was potently inhibited by *B. safensis* culture supernatants and concentrated peptide preparations, *Bacillus* species were not abundant in the surface microbiota of wooden boards in our study. We therefore examined the response of *L. monocytogenes* to a low dose of *B. safensis* culture supernatant (1.5%) that modestly inhibited *L. monocytogenes* growth (**Fig. S4**). Using a stringent fold change cut off, we found this treatment to significantly up-regulate of 25 genes by ≥4-fold, and significantly down-regulate 16 genes by ≥4-fold compared to untreated *L. monocytogenes* cultures (**Fig. 7**). Strikingly, all down-regulated genes map to the Φ10403S prophage or monocin locus (38), suggesting a role for prophage-encoded genes in antimicrobial stress. The up-regulated genes included a drug efflux system (*lmo2087*-*lmo2088*), consistent with an antimicrobial stress response. Furthermore, a phosphate transport system, homologous to the high-affinity PstSCAB phosphate transporter (39), was also up-regulated, suggesting that *B. safensis* supernatant causes phosphate starvation. Finally, an operon of unknown function, *lmo2567*-*lmo2568*, was also significantly up-regulated.

**Figure 7:**
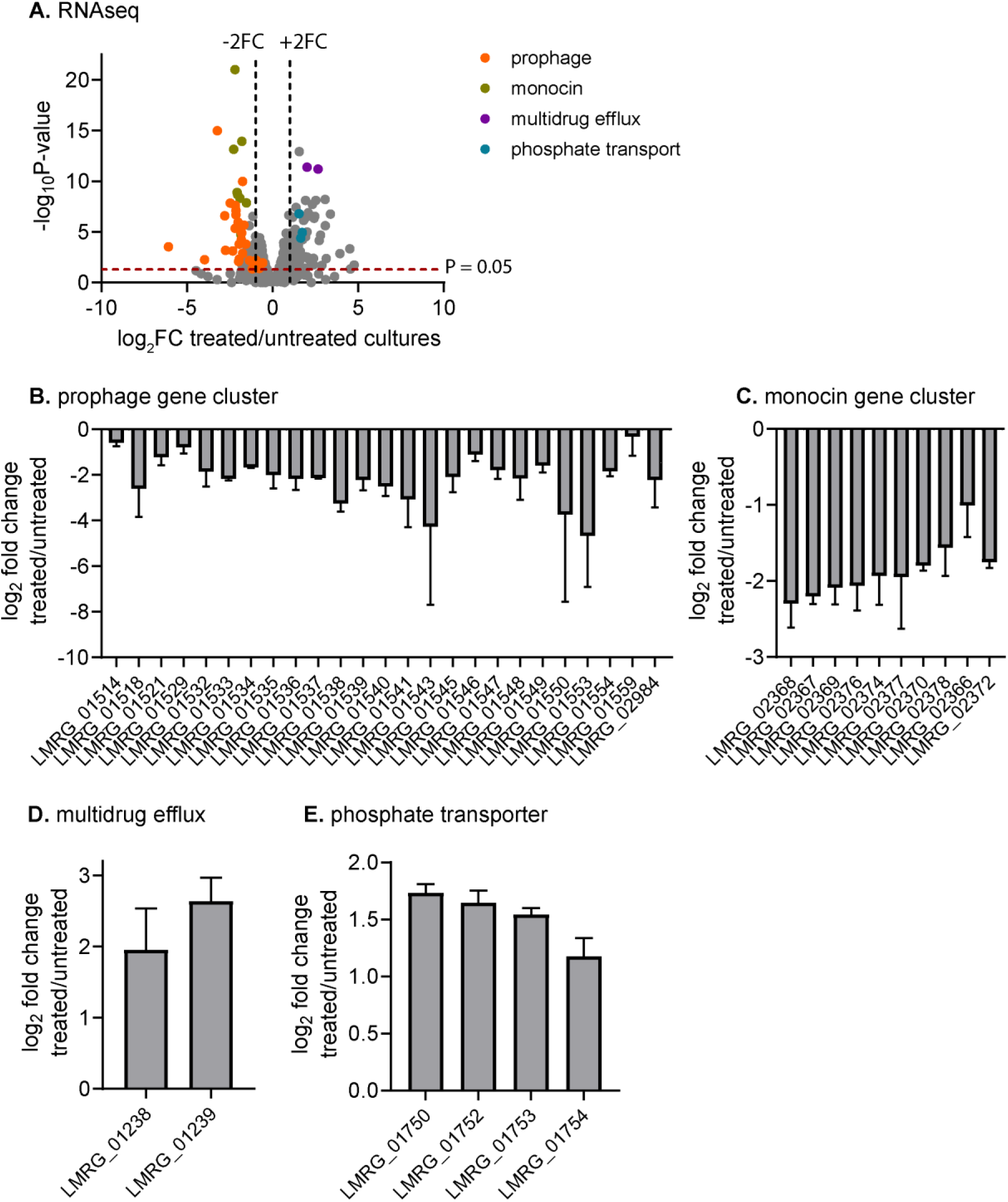
Transcriptional response of *L. monocytogenes* to *B. safensis* cell-free culture supernatant. A. Volcano plot depicts *L. monocytogenes* genes that were up- and down- regulated during growth in 1.5% *B. safensis* supernatant, compared with untreated cultures. **B-E.** Notable down- and up-regulated genes in treated vs. untreated *L. monocytogenes* cultures. Log_2_ fold change in gene expression was calculated by EdgeR, using normalized read counts per million.

## DISCUSSION

With a goal to expand our knowledge of wooden cheese board microbiota, we conducted a small survey of seven wooden boards from three different cheese makers in Wisconsin, United States. Our analysis confirms the diversity of bacterial communities on wooden boards in cheese ripening facilities. Of 93 genera identified on those boards, *Staphylococcus*, *Brevibacterium*, and *Brachybacterium* were universally present, and among most abundant bacteria in those microbial communities. On clean zones of wooden boards outside of cheese blocks, these three genera comprised up to 36%, 60%, and 32% of total bacterial abundance, respectively. Their abundance is congruent with two previous surveys of wooden boards in other locations (4, 5). Furthermore, we found these bacteria to also be prominent in board areas where cheese blocks had been placed, consistent with their ubiquitous presence on cheese rinds (21). All boards in our study also presented an abundance of *Nocardiopsis*, a genus of halotolerant bacteria that are also abundant in cheese rind communities, but not in other food products (21). Together, these observations suggest that *Staphylococcus*, *Brevibacterium*, and *Brachybacterium* are core members of wooden board microbial communities, and highlight the reciprocal microbial transfer between wood and cheese surfaces.

We reasoned that comparing the microbiota compositions of clean zones and “cheese” zones on native boards would reveal bacteria of wood and environmental origins. Indeed, principle component analysis indicates that bacterial communities on those zones were different on most boards. Upon further inspections, we observed a particularly high abundance of *Ruania* on clean zones of wooden boards from cheese maker 3 (up to 18% of bacterial communities), although their abundance was rather low in other cheese ripening facilities. Therefore, *Ruania* are likely associated with the ripening environment of cheese maker 3. With *Ruania* as an example, more thorough analyses are necessary to distinguish wood-associated bacteria from those of cheese origins, and their relative contributions to the ripening process.

As expected, washed and native boards in our study harbored distinguishable bacterial communities, and our data suggest that washing significantly reduces the abundance of *Staphylococcus* and increases the abundance of *Virgibacillus*. Larger surveys are necessary to establish the effects of sanitation on wooden board microbiota composition.

Our study also sought to assess *L. monocytogenes* survival on wooden cheese boards, with three technical improvements compared to a previous study (18). First, we applied grinding, a disruptive method, to maximize *L. monocytogenes* recovery from wood following surface inoculation (40). Second, recognizing that even the most effective method only recovers less than 30% of inoculated bacteria on wood, we normalized *L. monocytogenes* burdens to native board bacteria recovered from wooden boards. Raw *L. monocytogenes* counts on each board suggested that *L. monocytogenes* was inhibited. However, normalized burdens revealed one instance of *L. monocytogenes* replication, by 2-log of cfu in the “cheese” zone of board A. Therefore, internal standards are necessary controls for quantification of pathogens from wood. Finally, our study tracked *L. monocytogenes* burdens for 20 days following surface inoculation, much longer than a previous study (18), thereby providing safety data for longer storage of wooden boards.

Although *L. monocytogenes* grew within the cheese zone of board A, our aggregated data suggest that board wood inhibits *L. monocytogenes*. First, *L. monocytogenes* burdens declined in the clean area of board A, outside of the cheese block. Second, a reduction in *L. monocytogenes* burdens, by 0.5-2-log cfu, also occurred on other boards, both in the cheese and clean zones. Of note, board A was used for ripening of a soft cheese, which might explain its permissiveness for *L. monocytogenes* growth where there was leftover cheese. Combining the stability of native bacterial populations and the overall decline in *L. monocytogenes* counts, our data suggest that wooden boards are resilient to pathogen contamination over long-term storage. Analyses of additional boards over extended periods will be necessary to identify the core antimicrobial communities on wooden boards.

The inhibition of *L. monocytogenes* on wooden boards can be attributed to antimicrobial activities of both wood and the resident microbiota. Trees naturally produce antimicrobial compounds as defense against pathogens, and essential oils or polyphenolic compounds from wood have been demonstrated to effectively kill foodborne pathogens (19). However, since those studies mainly examined pathogen survival in highly concentrated antimicrobial extracts, the extent and kinetics of pathogen killing on cheese board woods remain to be determined.

Within wooden board microbiota, several prominent bacteria are known to produce antimicrobial compounds. For instance, *L. monocytogenes* is inhibited by various bacteriocins from lactic acid bacteria (23), phenazines and antimicrobial peptides from *Brevibacterium linens* (41, 42), and micrococcin P1 from *Staphylococcus equorum* (24). By systematically screening culturable bacteria from wooden cheese boards, we identified additional antimicrobial-producing bacteria whose abundances were low, such as *Serratia marcescens* and *Bacillus* species.

We focused on characterizing a wooden board *B. safensis* isolate, CB375, as a novel candidate for antimicrobial discovery against *L. monocytogenes*. *B. safensis* is a soil-dwelling bacterium that can colonize plant roots and stimulate plant growth (43–45), and some *B. safensis* strains have potent antifungal activity (26). Here, we found that *B. safensis* CB375 inhibits *L. monocytogenes* via secreted peptides or proteins that can be easily obtained and concentrated in the culture supernatant. Our bioinformatics analysis detected 14 biosynthetic gene clusters that encode antimicrobial peptides within *B. safensis* CB375 genome. Among BGCs that closely match known compounds, plantazolicin has a narrow spectrum activity and is unlikely to inhibit *L. monocytogenes* (31). Similarly, sporulation killing factors lyse kin cells within a *Bacillus* species population as a survival strategy under carbon source starvation, and are unlikely to act against other species (36, 37). Bacillibactin is a siderophore that can help *Bacillus* species in competitions with other bacteria, such as *Pseudomonas* (30, 46), but its effect against *L. monocytogenes* has not been tested. Bacilysin from *B. subtilis* is important for cellular differentiation and can inhibit other bacteria, such as *Campylobacter jejuni* (34, 47), but its antimicrobial activity has not been broadly characterized. The remaining 10 BGCs in *B. safensis* CB375 genome have low matches with known BGCs, or are unique to *B. safensis*. Purification and characterization of their products will be a promising direction in antimicrobial discovery.

*L. monocytogenes* strain 10403S encodes two phage-like elements, a Φ10403S prophage and a monocin that is predicted to attack other bacteria (38). Both elements were significantly down-regulated in response to a sub-inhibitory concentration of *B. safensis* supernatant. As for other bacteria, prophage and monocin gene expression in *L. monocytogenes* is induced by SOS response, activated upon treatments with DNA damaging reagents (38). The expression of these loci is also inhibited by prophage-encoded AriS, which suppresses SOS response, and de-repressed by monocin-encoded MpaR (38, 48). Therefore, the repression of prophage and monocin suggests that *B. safensis*-derived antimicrobial peptides might activate AriS, inhibit MpaR, or suppress SOS response by other unknown mechanisms. Among up-regulated genes, an increased expression of drug efflux is consistent with an antimicrobial stress response. The mechanisms that activate the phosphate transporter operon *pstSCAB* are much less clear. In Gram-negative bacteria, the *pstSCAB* operon is part of the phosphate starvation response that is activated by the PhoPR two-component system (49). Although *L. monocytogenes* harbors a PhoR homolog (50), its expression was not induced concomitantly with *pstSCAB* by *B. safensis* supernatant, and the PhoR regulon is not yet determined. Identification of the stimuli that activate phosphate transport and its role in antimicrobial response will expand research avenues for *L. monocytogenes* inhibition.

In summary, our study expands our knowledge of the microbial ecology on wooden boards used in cheese ripening. Our findings indicate an inhibitory effect of clean wooden boards against surface-inoculated *L. monocytogenes*. The relative contributions of wood and the resident microbiota to pathogen inhibition are unknown, but our identification of several bacteria that inhibit *L. monocytogenes* highlight wooden board microbiota as a source for antimicrobial discovery.

## MATERIALS AND METHODS

### Extraction of genomic DNA and sequencing of bacterial amplicons

Genomic DNA (gDNA) extraction of microbial community on the wooden board surfaces was performed based on a published procedure (4). Wood dust, obtained by drilling and shaving, was thoroughly resuspended in phosphate-buffered saline by gentle rotation, pelleted by centrifugation, and mechanically disrupted by bead beating, with the addition of phenol and 20% sodium dodecyl sulfate. Lysates were then centrifuged, and DNA was extracted and purified using phenol:chloroform:isoamyl alcohol (25:24:1, pH 8) (Thermo Fisher Scientific), precipitated, and washed with ethanol. DNA samples were quantified with a Qubit fluorometer (Invitrogen).

Total genomic DNA samples were PCR amplified for the 16S rRNA V4 region using universal primers as previously published (51). Briefly, 50 ng of DNA was used as template in each reaction, amplified with HotStart Ready Mix (KAPA Biosystems). PCR products were gel purified, quantified with a Qubit fluorometer, pooled in equimolar ratio, and sequenced on an Illumina MiSeq using a v2 kit to generate paired ended 250 bp reads with custom sequencing primers.

### Sequence processing and analysis

Sequences were demultiplexed on the Illumina MiSeq system and quality filtered using q2- demux plugin, followed by denoising with DADA2 (52). Subsequent processing was performed using QIIME 2 (53). Amplicon sequence variants were aligned using mafft (via q2-alignment) (54) and used to construct a phylogeny with fasttree2 (via q2-phylogeny) (55). Taxonomy was assigned to ASVs using the q2-feature-classifier. Relative abundances of top 20 taxa among all the samples were plotted using phyloseq and fantaxtic R packages (56, 57).

### Survival of *L. monocytogenes* on wooden board surface

Wooden boards were obtained from three cheese makers in Wisconsin (United States), cut into 24 blocks of 2 x 2 inches or 3 x 3 inches. Blocks from clean areas were separated from those inside the round area where cheese had been placed during ripening. *L. monocytogenes* 10403S (streptomycin resistant) was grown in Brain Heart Infusion broth, washed, resuspended in PBS, and spread evenly on each wooden board block. The blocks were placed in a closed container, maintained at 85% relative humidity using a saturated K_2_SO_4_ solution. All blocks were incubated at 12°C and relative humidity was monitored using a digital hygrometer. For *L. monocytogenes* recovery at each time point, wooden blocks were shaved off the surface and ground into small bits inside a clean beaker. Bacterial counts on day 0 were obtained 30 minutes after surface inoculation to allow *L. monocytogenes* impregnation on wood. Wood shavings and bits were thoroughly resuspended in PBS, with gentle disruption using sterile glass beads. Serial dilutions were plated on BHI with 200 µg streptomycin to count for *L. monocytogenes* cfu, or BHI without antibiotics for total bacteria. Native board-associated bacterial counts were calculated as total bacterial counts subtracted by *L. monocytogenes* counts.

### Isolation and identification of wooden board bacteria that inhibit *L. monocytogenes*

Wooden boards were shaved and ground to obtain wood bits, which were thoroughly resuspended in PBS by gentle vortexing. The PBS slurries were plated onto the following agar media: BHI, TSB (Trypic Soy Broth), PCAMS (22), and MRS. BHI, TSB, and PCAMS agar were incubated aerobically at 30°C. MRS agar was incubated at 37°C in a hypoxic chamber (4% CO_2_). Bacterial isolates were purified based on colony morphologies, and spotted onto a lawn of 10^5^ cfu *L. monocytogenes*. Bacterial isolates that produced zones of clearance were purified and tested for at least four more rounds. Initial identification was achieved using a MALDI-TOF Bruker Biotyper at the Wisconsin Veterinary Diagnostic Laboratory. For further identification, bacterial DNA was PCR-amplified for the V4-V9 regions using primers 515F and 1492R as previously published (58). Bacterial species were identified using BLAST, with sequencing reads as the query.

### Whole genome sequencing and genomic analyses

Genomic DNA was extracted from *Bacillus safensis* using phenol-chloroform (pH 8.0), precipitated and washed with ethanol, and resuspended in water. Illumina whole genome sequencing was performed at SeqCenter (Pittsburgh, USA) to obtain 2x150 bp reads at the depth of 1.33 million reads per sample. Quality control and adapter trimming were performed with fastqc. Contig and genome assembly were performed de novo by SPAdes (59), with quality assessment by QUAST (60). BV-BRC web resources were used to confirm taxonomic identification, find similar genomes (abou t 100 genomes), and constructed a phylogenetic tree. *B. safensis* CB375 and closely related genomes, revealed by the phylogenetic tree, were calculated for average nucleotide identity using KBase. Genome was annotated using Prokka (61), antimicrobial resistance genes were identified by CARD (29), prophages were identified by Phigaro and VirSorter2 (62, 63). Genome map was generated with Proksee (64). Biosynthetic gene clusters (BGCs) were predicted using antiSMASH 6.0 (65) and zol (66).

### Inhibition and killing of *L. monocytogenes* by *B. safensis*

For inhibition on BHI agar, *B. safensis* was streaked on one half of petri dishes and pre-incubated for 0-4 days prior to spotting of serial dilutions of a *L. monocytogenes* culture in the other half of the petri dish. BHI agar was further incubated for another 16 hours at 37°C to assess *L. monocytogenes* growth.

Cell-free culture supernatants were obtained from *B. safensis* cultures grown in BHI broth at 30°C for 16-20 hours and filtered through a 0.22 µm PES membrane. As a control, cell-free culture supernatants were also obtained from *L. monocytogenes* cultures grown at 37°C in BHI broth. Inhibition of *L. monocytogenes* in BHI broth was assessed in 96-well plates containing BHI only, or BHI containing indicated concentrations of cell-free culture supernatants. Bacterial growth was measured by OD_600_ at 37°C with intermittent shaking for 14 hours in a plate reader. For each strain, growth rates at each supernatant concentration were normalized to growth rate in BHI only of the same strain.

Secreted proteins and peptides from *B. safensis* were concentrated by adding a saturated ammonium sulfate solution (at a final 80%) to cell-free culture supernatants. Precipitates were separated by centrifugation and resuspended in PBS. Protein concentration was quantified by Bradford assay (Bio-Rad), using bovine serum albumin as standards. For proteinase K treatment, proteinase K was added to concentrated protein/peptide preparations at 1 mg/mL final concentration, incubated at 37°C for 2 hours, and 0.2mM PMSF was added to inactivate proteinase K. As a control, 0.2mM PMSF was added to another aliquot of concentrated proteins/peptides without proteinase K. For mock treatment, protein preparations were fincubated at 37°C for 2 hours.

For assessment of *L. monocytogenes* killing, concentrated protein preparations from *B. safensis* culture supernatant were diluted to 4 µg/mL, and added to *L. monocytogenes* cell suspensions in PBS. *L. monocytogenes* cfu were assessed at 0 and 1 hour post treatment, by plating serial dilutions on BHI agar + 200 µg/mL streptomycin.

### RNAseq and data analysis

*L. monocyotogenes* was grown in BHI to mid-log phase (OD_600_ ∼0.5) without or with 1.5% *B. safensis* cell-free culture supernatant. RNA was extracted and contaminated DNA was removed as previously described. RNAseq was performed at SeqCenter (Pittsburg, PA) to achieve 12 million 2 x 150 bp reads per sample. Reads were mapped to the *L. monocytogenes* 10403S genome (NCBI:txid393133). Differentially expressed gene analysis was performed by EdgeR (67).

## REFERENCES

1. Lortal S, Licitra G, Valence F. 2014. Wooden Tools: Reservoirs of Microbial Biodiversity in Traditional Cheesemaking. Microbiol Spectr 2:CM-0008–2012.

2. Mallia S, Carpino S, Corallo L, Tuminello L, Gelsomino R, Licitra G. 2005. Effects of Aroma Profiles of Piacentinu and Ricotta Cheese Using Different Tool Materials during Cheesemaking. Special Publication - Royal Society of Chemistry 300:23.

3. Licitra G, Caccamo M, Valence F, Lortal S. 2017. Traditional wooden equipment used for cheesemaking and their effect on quality, p.. *In* Papademas, P, Bintsis, T (eds.), Global Cheesemaking Technology: Cheese Quality and Characteristics. John Wiley & sons, Inc.

4. Wadhawan K, Steinberger AJ, Rankin SA, Suen G, Czuprynski CJ. 2021. Characterizing the microbiota of wooden boards used for cheese ripening. JDS Communications 2:171–176.

5. Settanni L, Busetta G, Puccio V, Licitra G, Franciosi E, Botta L, Di Gerlando R, Todaro M, Gaglio R. 2021. In-Depth Investigation of the Safety of Wooden Shelves Used for Traditional Cheese Ripening. Appl Environ Microbiol 87.

6. Mariani C, Briandet R, Chamba J-F, Notz E, Carnet-Pantiez A, Eyoug RN, Oulahal N. 2007. Biofilm ecology of wooden shelves used in ripening the French raw milk smear cheese Reblochon de Savoie. J Dairy Sci 90:1653–1661.

7. Irlinger F, Layec S, Hélinck S, Dugat-Bony E. 2015. Cheese rind microbial communities: diversity, composition and origin. FEMS Microbiol Lett 362:1–11.

8. Maury MM, Bracq-Dieye H, Huang L, Vales G, Lavina M, Thouvenot P, Disson O, Leclercq A, Brisse S, Lecuit M. 2019. Hypervirulent Listeria monocytogenes clones’ adaption to mammalian gut accounts for their association with dairy products. Nat Commun 10:2488.

9. Nightingale KK, Schukken YH, Nightingale CR, Fortes ED, Ho AJ, Her Z, Grohn YT, McDonough PL, Wiedmann M. 2004. Ecology and transmission of Listeria monocytogenes infecting ruminants and in the farm environment. Appl Environ Microbiol 70:4458–4467.

10. Sonnier JL, Karns JS, Lombard JE, Kopral CA, Haley BJ, Kim S-W, Van Kessel JAS. 2018. Prevalence of Salmonella enterica, Listeria monocytogenes, and pathogenic Escherichia coli in bulk tank milk and milk filters from US dairy operations in the National Animal Health Monitoring System Dairy 2014 study. J Dairy Sci 101:1943–1956.

11. Haley BJ, Sonnier J, Schukken YH, Karns JS, Van Kessel JAS. 2015. Diversity of Listeria monocytogenes within a U.S. dairy herd, 2004-2010. Foodborne Pathog Dis 12:844–850.

12. Almeida G, Magalhães R, Carneiro L, Santos I, Silva J, Ferreira V, Hogg T, Teixeira P. 2013. Foci of contamination of Listeria monocytogenes in different cheese processing plants. Int J Food Microbiol 167:303–309.

13. Chowdhury B, Anand S. 2023. Environmental persistence of *Listeria monocytogenes* and its implications in dairy processing plants. Compr Rev Food Sci Food Saf 22:4573–4599.

14. Guldimann C, Boor KJ, Wiedmann M, Guariglia-Oropeza V. 2016. Resilience in the Face of Uncertainty: Sigma Factor B Fine-Tunes Gene Expression To Support Homeostasis in Gram-Positive Bacteria. Appl Environ Microbiol 82:4456–4469.

15. Colagiorgi A, Bruini I, Di Ciccio PA, Zanardi E, Ghidini S, Ianieri A. 2017. Listeria monocytogenes Biofilms in the Wonderland of Food Industry. Pathogens 6.

16. Valderrama WB, Cutter CN. 2013. An ecological perspective of Listeria monocytogenes biofilms in food processing facilities. Crit Rev Food Sci Nutr 53:801–817.

17. Ismail R, Aviat F, Gay-Perret P, Le Bayon I, Federighi M, Michel V. 2017. An assessment of L. monocytogenes transfer from wooden ripening shelves to cheeses: Comparison with glass and plastic surfaces. Food Control 73:273–280.

18. Mariani C, Oulahal N, Chamba J-F, Dubois-Brissonnet F, Notz E, Briandet R. 2011. Inhibition of Listeria monocytogenes by resident biofilms present on wooden shelves used for cheese ripening. Food Control 22:1357–1362.

19. Aviat F, Gerhards C, Rodriguez-Jerez J, Michel V, Bayon I Le, Ismail R, Federighi M. 2016. Microbial Safety of Wood in Contact with Food: A Review. Compr Rev Food Sci Food Saf 15:491– 505.

20. Wadhawan K, Steinberger A, Rankin S, Suen G, Czuprynski C. 2023. Inhibition of Listeria monocytogenes by Broth Cultures of Surface Microbiota of Wooden Boards Used in Cheese Ripening. Applied Sciences 13:5872.

21. Wolfe BE, Button JE, Santarelli M, Dutton RJ. 2014. Cheese rind communities provide tractable systems for in situ and in vitro studies of microbial diversity. Cell 158:422–433.

22. Cosetta CM, Wolfe BE. 2020. Deconstructing and Reconstructing Cheese Rind Microbiomes for Experiments in Microbial Ecology and Evolution. Curr Protoc Microbiol 56:e95.

23. Sugrue I, Ross RP, Hill C. 2024. Bacteriocin diversity, function, discovery and application as antimicrobials. Nat Rev Microbiol 22:556–571.

24. Carnio MC, Höltzel A, Rudolf M, Henle T, Jung G, Scherer S. 2000. The Macrocyclic Peptide Antibiotic Micrococcin P _1_ Is Secreted by the Food-Borne Bacterium *Staphylococcus equorum* WS 2733 and Inhibits *Listeria monocytogenes* on Soft Cheese. Appl Environ Microbiol 66:2378–2384.

25. Decker T, Rautenbach M, Khan S, Khan W. 2024. Antibacterial efficacy and membrane mechanism of action of the *Serratia* -derived non-ionic lipopeptide, serrawettin W2-FL10. Microbiol Spectr 12.

26. Mayer FL, Kronstad JW. 2017. Disarming Fungal Pathogens: *Bacillus safensis* Inhibits Virulence Factor Production and Biofilm Formation by *Cryptococcus neoformans* and *Candida albicans*. mBio 8.

27. Ragon M, Wirth T, Hollandt F, Lavenir R, Lecuit M, Le Monnier A, Brisse S. 2008. A new perspective on Listeria monocytogenes evolution. PLoS Pathog 4:e1000146.

28. Bécavin C, Bouchier C, Lechat P, Archambaud C, Creno S, Gouin E, Wu Z, Kühbacher A, Brisse S, Pucciarelli MG, García-del Portillo F, Hain T, Portnoy DA, Chakraborty T, Lecuit M, Pizarro-Cerdá J, Moszer I, Bierne H, Cossart P. 2014. Comparison of widely used Listeria monocytogenes strains EGD, 10403S, and EGD-e highlights genomic variations underlying differences in pathogenicity. mBio 5:e00969–14.

29. Alcock BP, Huynh W, Chalil R, Smith KW, Raphenya AR, Wlodarski MA, Edalatmand A, Petkau A, Syed SA, Tsang KK, Baker SJC, Dave M, McCarthy MC, Mukiri KM, Nasir JA, Golbon B, Imtiaz H, Jiang X, Kaur K, Kwong M, Liang ZC, Niu KC, Shan P, Yang JYJ, Gray KL, Hoad GR, Jia B, Bhando T, Carfrae LA, Farha MA, French S, Gordzevich R, Rachwalski K, Tu MM, Bordeleau E, Dooley D, Griffiths E, Zubyk HL, Brown ED, Maguire F, Beiko RG, Hsiao WWL, Brinkman FSL, Van Domselaar G, McArthur AG. 2023. CARD 2023: expanded curation, support for machine learning, and resistome prediction at the Comprehensive Antibiotic Resistance Database. Nucleic Acids Res 51:D690–D699.

30. Dimopoulou A, Theologidis I, Benaki D, Koukounia M, Zervakou A, Tzima A, Diallinas G, Hatzinikolaou DG, Skandalis N. 2021. Direct Antibiotic Activity of Bacillibactin Broadens the Biocontrol Range of Bacillus amyloliquefaciens MBI600. mSphere 6.

31. Molohon KJ, Saint-Vincent PMB, Park S, Doroghazi JR, Maxson T, Hershfield JR, Flatt KM, Schroeder NE, Ha T, Mitchell DA. 2016. Plantazolicin Is an Ultranarrow-Spectrum Antibiotic That Targets the *Bacillus anthracis* Membrane. ACS Infect Dis 2:207–220.

32. González-Pastor JE, Hobbs EC, Losick R. 2003. Cannibalism by Sporulating Bacteria. Science (1979) 301:510–513.

33. Höfler C, Heckmann J, Fritsch A, Popp P, Gebhard S, Fritz G, Mascher T. 2016. Cannibalism stress response in Bacillus subtilis. Microbiology (N Y) 162:164–176.

34. Erega A, Stefanic P, Danevčič T, Smole Možina S, Mandic Mulec I. 2022. Impact of *Bacillus subtilis* Antibiotic Bacilysin and *Campylobacter jejuni* Efflux Pumps on Pathogen Survival in Mixed Biofilms. Microbiol Spectr 10.

35. Han X, Shen D, Xiong Q, Bao B, Zhang W, Dai T, Zhao Y, Borriss R, Fan B. 2021. The Plant-Beneficial Rhizobacterium Bacillus velezensis FZB42 Controls the Soybean Pathogen Phytophthora sojae Due to Bacilysin Production. Appl Environ Microbiol 87.

36. Zhang L, Sun C. 2018. Fengycins, Cyclic Lipopeptides from Marine Bacillus subtilis Strains, Kill the Plant-Pathogenic Fungus Magnaporthe grisea by Inducing Reactive Oxygen Species Production and Chromatin Condensation. Appl Environ Microbiol 84.

37. Akers HA. 1983. Isolation of the Siderophore Schizokinen from Soil of Rice Fields. Appl Environ Microbiol 45:1704–1706.

38. Argov T, Sapir SR, Pasechnek A, Azulay G, Stadnyuk O, Rabinovich L, Sigal N, Borovok I, Herskovits AA. 2019. Coordination of cohabiting phage elements supports bacteria–phage cooperation. Nat Commun 10:5288.

39. Choi S, Jeong G, Choi E, Lee E-J. 2022. A Dual Regulatory Role of the PhoU Protein in Salmonella Typhimurium. mBio 13.

40. Ismaïl R, Le Bayon I, Michel V, Jequel M, Kutnik M, Aviat F, Fédérighi M. 2015. Comparative Study of Three Methods for Recovering Microorganisms from Wooden Surfaces in the Food Industry. Food Anal Methods 8:1238–1247.

41. Motta AS, Brandelli A. 2002. Characterization of an antibacterial peptide produced by Brevibacterium linens. J Appl Microbiol 92:63–70.

42. Cumsille A, Serna-Cardona N, González V, Claverías F, Undabarrena A, Molina V, Salvà-Serra F, Moore ERB, Cámara B. 2023. Exploring the biosynthetic gene clusters in Brevibacterium: a comparative genomic analysis of diversity and distribution. BMC Genomics 24:622.

43. Mateus JR, Marques JM, Dal’Rio I, Vollú RE, Coelho MRR, Seldin L. 2019. Response of the microbial community associated with sweet potato (Ipomoea batatas) to Bacillus safensis and Bacillus velezensis strains. Antonie Van Leeuwenhoek 112:501–512.

44. Prakash J, Arora NK. 2020. Development of Bacillus safensis-based liquid bioformulation to augment growth, stevioside content, and nutrient uptake in Stevia rebaudiana. World J Microbiol Biotechnol 36:8.

45. Romero-Severson J, Moran TE, Shrader DG, Fields FR, Pandey-Joshi S, Thomas CL, Palmer EC, Shrout JD, Pfrender ME, Lee SW. 2021. A Seed-Endophytic Bacillus safensis Strain With Antimicrobial Activity Has Genes for Novel Bacteriocin-Like Antimicrobial Peptides. Front Microbiol 12.

46. Lyng M, Jørgensen JPB, Schostag MD, Jarmusch SA, Aguilar DKC, Lozano-Andrade CN, Kovács ÁT. 2024. Competition for iron shapes metabolic antagonism between *Bacillus subtilis* and *Pseudomonas marginalis*. ISME J 18.

47. Schoenborn AA, Yannarell SM, Wallace ED, Clapper H, Weinstein IC, Shank EA. 2021. Defining the Expression, Production, and Signaling Roles of Specialized Metabolites during Bacillus subtilis Differentiation. J Bacteriol 203.

48. Azulay G, Pasechnek A, Stadnyuk O, Ran-Sapir S, Fleisacher AM, Borovok I, Sigal N, Herskovits AA. 2022. A dual-function phage regulator controls the response of cohabiting phage elements via regulation of the bacterial SOS response. Cell Rep 39:110723.

49. Moreau PL. 2023. Regulation of phosphate starvation-specific responses in Escherichia coli. Microbiology (N Y) 169.

50. Alejandro-Navarreto X, Freitag NE. 2024. Revisiting old friends: updates on the role of two-component signaling systems in *Listeria monocytogenes* survival and pathogenesis. Infect Immun 92.

51. Dias J, Marcondes MI, Motta de Souza S, Cardoso da Mata E Silva B, Fontes Noronha M, Tassinari Resende R, Machado FS, Cuquetto Mantovani H, Dill-McFarland KA, Suen G. 2018. Bacterial Community Dynamics across the Gastrointestinal Tracts of Dairy Calves during Preweaning Development. Appl Environ Microbiol 84.

52. Callahan BJ, McMurdie PJ, Rosen MJ, Han AW, Johnson AJA, Holmes SP. 2016. DADA2: High-resolution sample inference from Illumina amplicon data. Nat Methods 13:581–583.

53. Bolyen E, Rideout JR, Dillon MR, Bokulich NA, Abnet CC, Al-Ghalith GA, Alexander H, Alm EJ, Arumugam M, Asnicar F, Bai Y, Bisanz JE, Bittinger K, Brejnrod A, Brislawn CJ, Brown CT, Callahan BJ, Caraballo-Rodríguez AM, Chase J, Cope EK, Da Silva R, Diener C, Dorrestein PC, Douglas GM, Durall DM, Duvallet C, Edwardson CF, Ernst M, Estaki M, Fouquier J, Gauglitz JM, Gibbons SM, Gibson DL, Gonzalez A, Gorlick K, Guo J, Hillmann B, Holmes S, Holste H, Huttenhower C, Huttley GA, Janssen S, Jarmusch AK, Jiang L, Kaehler BD, Kang K Bin, Keefe CR, Keim P, Kelley ST, Knights D, Koester I, Kosciolek T, Kreps J, Langille MGI, Lee J, Ley R, Liu Y-X, Loftfield E, Lozupone C, Maher M, Marotz C, Martin BD, McDonald D, McIver LJ, Melnik A V., Metcalf JL, Morgan SC, Morton JT, Naimey AT, Navas-Molina JA, Nothias LF, Orchanian SB, Pearson T, Peoples SL, Petras D, Preuss ML, Pruesse E, Rasmussen LB, Rivers A, Robeson MS, Rosenthal P, Segata N, Shaffer M, Shiffer A, Sinha R, Song SJ, Spear JR, Swafford AD, Thompson LR, Torres PJ, Trinh P, Tripathi A, Turnbaugh PJ, Ul-Hasan S, van der Hooft JJJ, Vargas F, Vázquez-Baeza Y, Vogtmann E, von Hippel M, Walters W, Wan Y, Wang M, Warren J, Weber KC, Williamson CHD, Willis AD, Xu ZZ, Zaneveld JR, Zhang Y, Zhu Q, Knight R, Caporaso JG. 2019. Reproducible, interactive, scalable and extensible microbiome data science using QIIME 2. Nat Biotechnol 37:852–857.

54. Katoh K. 2002. MAFFT: a novel method for rapid multiple sequence alignment based on fast Fourier transform. Nucleic Acids Res 30:3059–3066.

55. Price MN, Dehal PS, Arkin AP. 2010. FastTree 2 – Approximately Maximum-Likelihood Trees for Large Alignments. PLoS One 5:e9490.

56. Teunisse G. 2022. Fantaxtic - Nested Bar Plots for Phyloseq Data (Version 0.2.1).

57. McMurdie PJ, Holmes S. 2013. phyloseq: An R Package for Reproducible Interactive Analysis and Graphics of Microbiome Census Data. PLoS One 8:e61217.

58. Abellan-Schneyder I, Matchado MS, Reitmeier S, Sommer A, Sewald Z, Baumbach J, List M, Neuhaus K. 2021. Primer, Pipelines, Parameters: Issues in 16S rRNA Gene Sequencing. mSphere 6.

59. Prjibelski A, Antipov D, Meleshko D, Lapidus A, Korobeynikov A. 2020. Using SPAdes De Novo Assembler. Curr Protoc Bioinformatics 70.

60. Gurevich A, Saveliev V, Vyahhi N, Tesler G. 2013. QUAST: quality assessment tool for genome assemblies. Bioinformatics 29:1072–1075.

61. Seemann T. 2014. Prokka: rapid prokaryotic genome annotation. Bioinformatics 30:2068–2069.

62. Guo J, Bolduc B, Zayed AA, Varsani A, Dominguez-Huerta G, Delmont TO, Pratama AA, Gazitúa MC, Vik D, Sullivan MB, Roux S. 2021. VirSorter2: a multi-classifier, expert-guided approach to detect diverse DNA and RNA viruses. Microbiome 9:37.

63. Starikova E V, Tikhonova PO, Prianichnikov NA, Rands CM, Zdobnov EM, Ilina EN, Govorun VM. 2020. Phigaro: high-throughput prophage sequence annotation. Bioinformatics 36:3882–3884.

64. Grant JR, Enns E, Marinier E, Mandal A, Herman EK, Chen C, Graham M, Van Domselaar G, Stothard P. 2023. Proksee: in-depth characterization and visualization of bacterial genomes. Nucleic Acids Res 51:W484–W492.

65. Blin K, Shaw S, Kloosterman AM, Charlop-Powers Z, van Wezel GP, Medema MH, Weber T. 2021. antiSMASH 6.0: improving cluster detection and comparison capabilities. Nucleic Acids Res 49:W29–W35.

66. Salamzade R, Tran PQ, Martin C, Manson AL, Gilmore MS, Earl AM, Anantharaman K, Kalan L. 2023. zol & fai: large-scale targeted detection and evolutionary investigation of gene clusters 10.1101/2023.06.07.544063.

67. Robinson MD, McCarthy DJ, Smyth GK. 2010. <tt>edgeR</tt> : a Bioconductor package for differential expression analysis of digital gene expression data. Bioinformatics 26:139–140.

